# Mapping the Structural and Dynamic Determinants of pH-sensitive Heparin Binding to Granulocyte Macrophage-colony Stimulating Factor

**DOI:** 10.1101/2020.03.11.987370

**Authors:** Jennifer Y. Cui, Fuming Zhang, Robert J. Linhardt, George P. Lisi

**Affiliations:** Department of Molecular Biology, Cell Biology & Biochemistry, Brown University, Providence, RI 02903 USA; Departments of Chemistry, Biology, & Chemical & Biological Engineering, Rensselaer Polytechnic Institute, Troy, NY 12180 USA

**Keywords:** heparin, GMCSF, NMR, SPR, protein dynamics

## Abstract

GMCSF is an immunomodulatory cytokine that is harnessed as a therapeutic. GMCSF is known to interact with other clinically important molecules, such as heparin, suggesting that endogenous and administered GMCSF has the potential to modulate orthogonal treatment outcomes. Thus, molecular level characterization of GMCSF and its interactions with other biologically active compounds is critical to understanding these mechanisms and predicting clinical outcomes. Here, we dissect the molecular motions and structural contributions that facilitate the GMCSF-heparin interaction, previously shown to be pH-dependent, using NMR spectroscopy, SPR, and molecular docking. We find that GMCSF and heparin binding is related to a change in flexibility reflected in the dynamic profile of GMCSF at acidic pH. The molecular motions driving this interaction largely occur on the ms-µs timescale. Interestingly, we find that GMCSF and heparin binding is not only pH-dependent but is also heparin chain length dependent. We propose a mechanism where a positive binding pocket that is not fully solvent accessible at neutral pH becomes more accessible at acidic pH, allowing heparin to dock with the protein.

## Introduction

Granulocyte macrophage colony stimulating factor (GMCSF, or CSF2) is a mediator of cellular expansion and differentiation for myeloid progenitors in bone marrow. GMCSF plays an immunoregulatory role as a pathogenic pro-inflammatory agent in autoimmune diseases such as multiple sclerosis (MS) and rheumatoid arthritis (RA)^1,2^ but has also been utilized in a clinical setting as a therapeutic for immunocompromised patients to simulate recovery of mature white blood cell populations.^3^ Given that GMCSF has been shown to be both a protective and pathogenic signal transduction mediator in innate immune responses and has been employed as a therapeutic for immunosuppressed patients, molecules that potentially modulate GMCSF are of interest. The interactions of GMCSF with several biologically relevant molecules, including heparin, heparan sulfate, chondroitin sulfates, and nucleotide triphosphates have been investigated to various degrees at the biophysical level.^4-8^ Importantly, heparin was previously shown as an inhibitor of a nucleotide binding site that correlates directly to the proliferative ability of GMCSF.^9^

Heparan sulfate is a biologically important molecule for its endogenous function as a cell surface glycosaminoglycan (GAG) and it is structurally related to heparin, which is clinically used as an anticoagulant. Unfractionated heparin, derived from porcine intestinal mucosa and the most common form of heparin-based drug, is inexpensive and has a known antidote, protamine sulfate.^10,11^ However, unfractionated heparin is also associated with severe adverse immune reactions in a small fraction of patients, including the development of heparin-induced thrombocytopenia (HIT).^12^ HIT occurs when heparin binds to the chemokine platelet factor 4 (PF4), causing it to undergo a conformational change resulting in an immunological response against the resulting ultra-large complexes.^13,14^ Low molecular weight heparins, developed as alternative therapeutics to unfractionated heparin, show reduced frequencies of adverse immune reactions.^15^ Despite the attenuated instances of HIT in patients, the clinical outcomes of HIT are still severe and often life threatening.

Previous work has shown that GMCSF can interact with heparin at acidic pH (between pH 4-5), but does not interact at neutral pH.^4,7,8^ The interaction of GMCSF with negatively charged heparin molecules is reportedly dependent on the presence of three histidine residues (His15, His83, and His87) that become positively charged at acidic pH.^7^ An additional residue, Lys85, is also thought to be involved. Interestingly, the clinical administration of GMCSF and heparin often occur in patients that are experiencing severe pathologies with effects that alter local physiological pH balance, such as asthmatic conditions, sepsis, and cancer.^16-18^ Given the frequency of unfractionated and low molecular weight heparin use in clinical settings, and the role of GMCSF in altering immune responses, it is important to elucidate the nuanced interactions between these molecules, both of which exist endogenously and are administered as independent therapeutics, creating ample opportunity for them to exist in the same physiological space.

We explored the structural, dynamic, and biophysical properties of these complexes to better understand the interactions between GMCSF and heparin. It is well known that structural dynamics can reveal mechanistic details of protein-ligand interactions, and although a binding site for heparin on GMCSF has been proposed, there is no molecular level information about this interaction. In this study, we use surface plasmon resonance (SPR) and solution nuclear magnetic resonance (NMR) techniques to update known binding interactions of unfractionated heparin with GMCSF, as well as provide novel probes of low molecular weight heparin-GMCSF interactions. Additionally, we describe the structural and dynamical changes in the GMCSF protein at acidic pH and in the presence of low molecular weight heparins at neutral and acidic pH.

## Results

### GMCSF-heparin interactions are size and pH dependent

In order to assess the binding of unfractionated heparin and heparin oligosaccharides (oligos, **Figure 1a**) to GMCSF (**Figure 1b**), we used SPR. Sensorgrams collected with varying protein concentrations were globally fitted with 1:1 Langmuir model, and the binding kinetics and affinities are presented in **Table 1**.

**Table 1.** pH-dependent binding affinities of GMCSF for heparin.

| Interaction | $k_a$ (1/s) | $k_d$ (1/s) | $K_D$ (M) |
| --- | --- | --- | --- |
| pH 7.4 | $6.3 \times 10^{-2}$<br>($\pm 3.0 \times 10^{-3}$) | $4.2 \times 10^{-4}$<br>($\pm 2.7 \times 10^{-5}$) | $6.8 \times 10^{-6}$ |
| pH 5.5 | $6.8 \times 10^{-2}$<br>($\pm 2.0 \times 10^{-3}$) | $3.9 \times 10^{-4}$<br>($\pm 1.2 \times 10^{-5}$) | $5.8 \times 10^{-6}$ |
| pH 4.5 | 2.8<br>($\pm 9.0 \times 10^{-2}$) | $5.6 \times 10^{-4}$<br>($\pm 2.0 \times 10^{-5}$) | $2.0 \times 10^{-7}$ |

Sensorgrams of GMCSF-heparin interactions at pH 7.4, 5.5, and 4.5 show that the affinity of heparin for GMCSF increases with solution acidity, as shown by the reduction in *K*_*D*_ (**Figure 1c, Table 1**), which was particularly pronounced as the pH changed from 5.5 to 4.5 and is primarily the result of the increased association rate (k_a_) of heparin to GMCSF.

**Figure 1.**
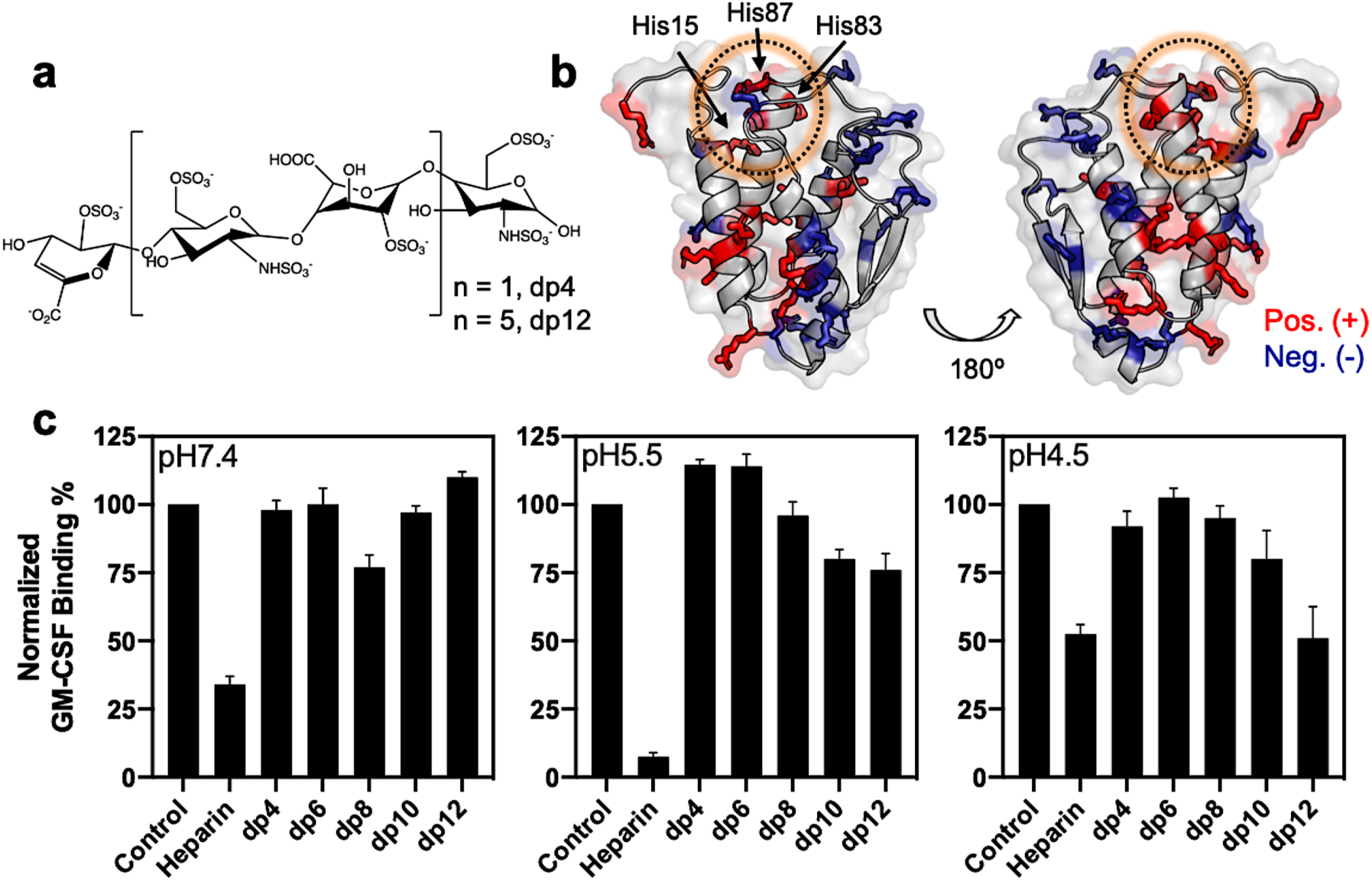
GMCSF-heparin interactions studied by SPR. **a**. Structure of the heparin oligosaccharide and major repeating sequence of heparin (brackets) with degree of polymerization (dp)4 and dp12. **b**. X-ray crystal structure of GMCSF (PDB 2GMF) with the distribution of positively charged residues (Arg, Lys; red) and negatively charged residues (Glu, Asp; blue) shown in sticks. The critical His residues clustered at the GMCSF termini are highlighted and circled. **c**. Normalized competitive binding between immobilized heparin and aqueous solutions of GMCSF (1 µM, no competing ligand) and GMCSF complexed with unfractionated heparin (negative and positive controls) or heparin oligosaccharide dp4, dp6, dp8, dp10, dp12 from SPR experiments conducted at pH 7.4, 5.5 and 4.5. Error bars were determined from replicate experiments.

We next performed solution/surface competition experiments where preformed complexes of GMCSF and variably sized heparin oligos (**Figure 1a**, dp4, dp6, dp8, dp10, dp12) were flowed over a chip with immobilized heparin at pH 5.5 to examine the effect of heparin oligo chain size on the GMCSF-heparin interaction. Complexes of GMCSF and heparin oligos that had impaired binding to the chip surface were considered to have stronger binding interactions. GMCSF with and without heparin were used as controls. At pH 5.5, no competitive interaction was observed when 1 mM dp4, dp6 or dp8 were pre-incubated with GMCSF (**Figure 1c**). Indeed, it appears that dp4 and dp6 may even slightly enhance GMCSF binding to immobilized heparin, relative to the control sample. Binding between surface-bound heparin and GMCSF was significantly diminished only in the presence of the dp10 and dp12 oligos and 1 mM soluble heparin completely blocked binding. Similar experiments were conducted at pH 7.4 and pH 4.5 suggesting that the interaction between the GMCSF and heparin is pH dependent and chain-length dependent beginning with a chain length of > 8. Based on these data, we used dp4 and dp12 heparin oligos as representatives for each of the observed binding characteristics of heparin in subsequent biophysical experiments.

### The GMCSF structure and molecular motions are altered by acidic pH

We obtained ^1^H^15^N heteronuclear single quantum coherence (HSQC) NMR spectra of GMCSF at pH 7.4 and pH 5.5 to investigate the effect of acidic conditions, which favor heparin binding, on the GMCSF structure and conformational dynamics (**Supplementary Figure 2**). We did not collect NMR data at pH 4.5, which crosses the isoelectric point of GMCSF, due to accelerated precipitation of the protein at NMR concentrations. Large chemical shift perturbations (Δδ) are observed at pH 5.5, highlighting changes in the local environments of GMCSF residues (**Figure 2a**), primarily clustered around the histidine triad on helices A and C, the loop connecting helix A and sheet β1, as well as the loop connecting sheet β1 and helix B. Longitudinal (*R*_1_) and transverse (*R*_2_) relaxation rates measured by NMR highlight several regions of multi-timescale conformational flexibility in GMCSF (**Figure 2b**,**c**). Here, the pH-dependent relaxation rates are plotted as the *R*_1_*R*_2_ product to suppress contributions of anisotropic molecular tumbling that complicate interpretations of chemical exchange.

**Figure 2.**
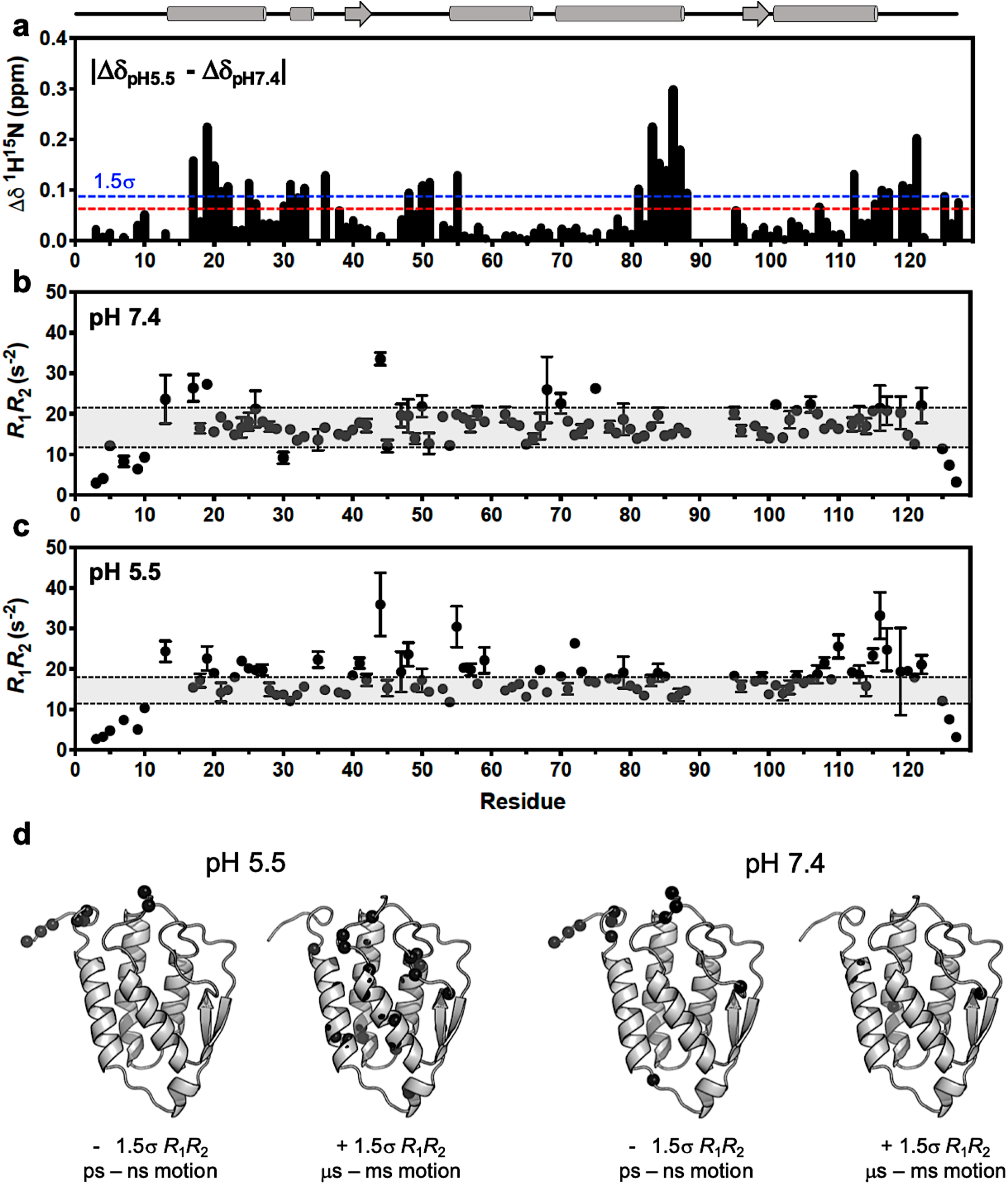
pH-dependent structural and dynamic changes in GMCSF. **a**. Combined 1H^15^N chemical shift perturbations (|Δδ_pH5.5_ - Δδ_pH7.4_|) calculated as 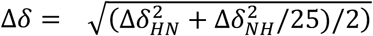. The red and blue lines denote the 10% trimmed mean and 1.5σ above the mean, respectively. A cartoon representation of the GMCSF structure is shown above the panel. **b**. NMR *R*_1_*R*_2_ relaxation parameters for apo GMCSF at pH 7.4. The gray shaded area denotes ± 1.5σ of the 10% trimmed mean of the data. **c**. NMR *R*_1_*R*_2_ relaxation parameters for apo GMCSF at pH 5.5. The gray shaded area denotes ± 1.5σ of the 10% trimmed mean of the data. **d**. Residues with *R*_1_*R*_2_ values below (−1.5σ) or above (+ 1.5σ) the 10% trimmed mean *R*_1_*R*_2_ are mapped onto the GMCSF structure (PDB 2GMF).

The average generalized order parameter was calculated with 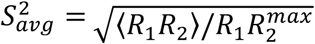, using values of <*R*_1_*R*_2_> and *R*_1_*R*_2_^max^ reflective of the entire data sets.^19^ At neutral pH, four residues in within the core of GMCSF (G75, N17, S44, V16, **Figure 2d**) display *R*_1_*R*_2_ values above 1.5σ of the 10% trimmed mean of all rates measured at pH 7.4 due to the influence of *R*_ex_ related to µs-ms motion. An additional ten residues show *R*_1_*R*_2_ values 1.5σ below the 10% trimmed mean, qualitatively suggesting ps-ns timescale motion, as the average generalized order parameter determined from the entire data set is *S*^2^ = 0.64. Unsurprisingly, the majority of residues with depressed *R*_1_*R*_2_ values are found at the N- and C-termini, while R30 and E45 are within loops or unstructured regions of the protein. Apo GMCSF is relatively inflexible near neutral pH, while at acidic pH, µs-ms motions are observed within the protein core and one face of the protein, involving 29 residues along helices A and C (**Figure 2d**). The contribution of fast timescale dynamics is essentially unchanged, restricted to terminal residues A3, R4, S5, S7, S9, T10, Q126, and E127 showing an average generalized order parameter of all residues of *S*^2^ = 0.69 at pH 5.5. The <*S*^2^> at pH 7.4 and pH 5.5 are comparable and suggest that the primary influence is largely dominated by µs-ms timescales.

### GMCSF structure and dynamics are altered in a pH- and heparin size-dependent manner

^15^N-labeled GMCSF was titrated with unfractionated heparin (containing low and high molecular weight species), dp4, and dp12 at both pH 7.4 and pH 5.5, to further characterize GMCSF and heparin binding. Statistically significant chemical shift perturbations (Δδ) were determined by averaging ^1^H^15^N combined Δδ values for all GMCSF-heparin samples at a given pH. At pH 7.4, unfractionated heparin did not cause any chemical shift perturbations in the GMCSF NMR spectrum (**Figure 3a**). Titration of GMCSF with dp4 at pH 7.4 perturbs only 2 resonances, corresponding to E14 and L55 on helices A/B **(Supplementary Figure 2a)**. Interestingly, when dp12 is titrated into GMCSF at pH 7.4, chemical shifts of 27 residues are significantly perturbed (**Figure 4b, Supplementary Figure 2b**).

**Figure 3.**
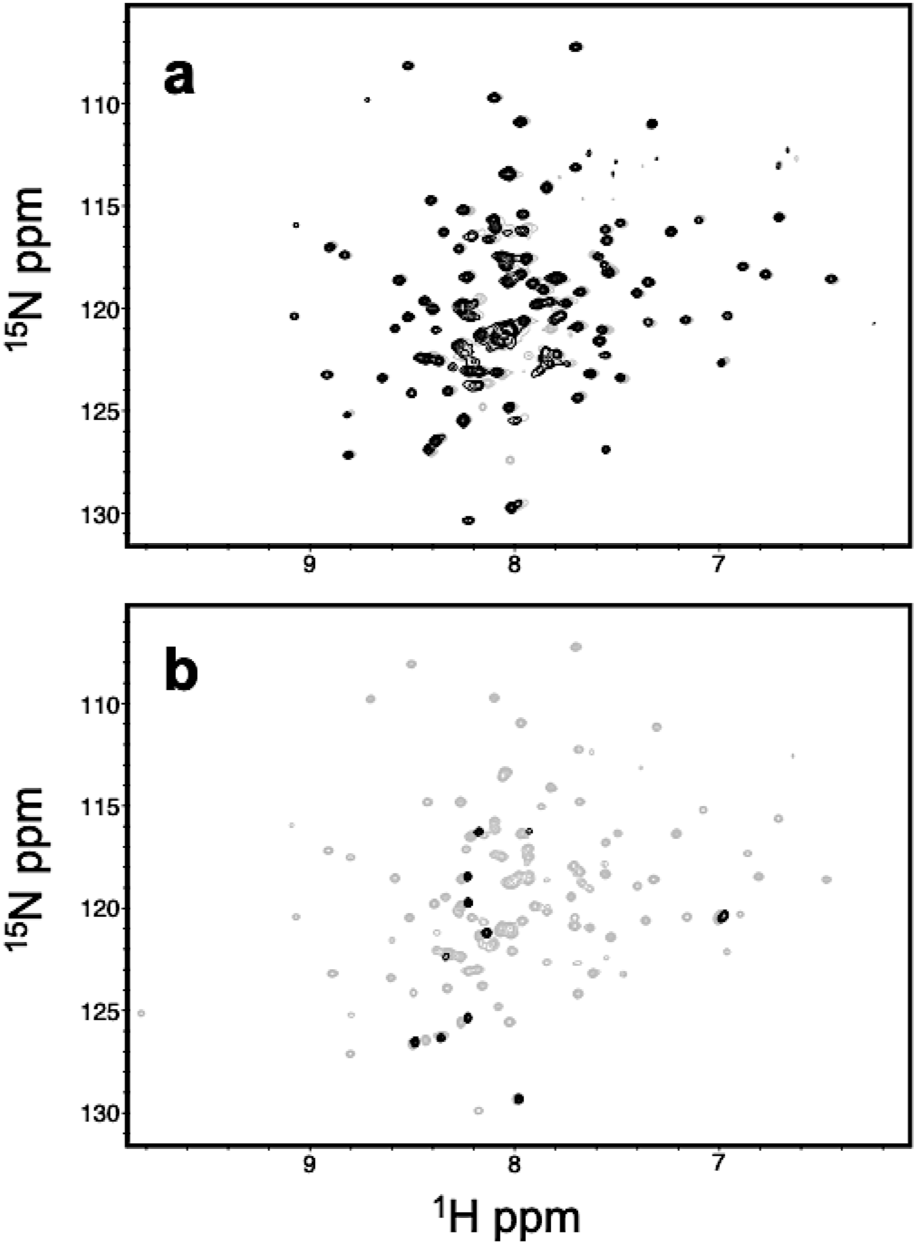
pH-dependent interaction of GMCSF with unfractionated heparin. **a**. ^1^H^15^N HSQC NMR spectrum of GMCSF at pH 7.4 (gray) and GMCSF saturated with unfractionated heparin (black). **b**. Identical experiment carried out at pH 5.5.

**Figure 4.**
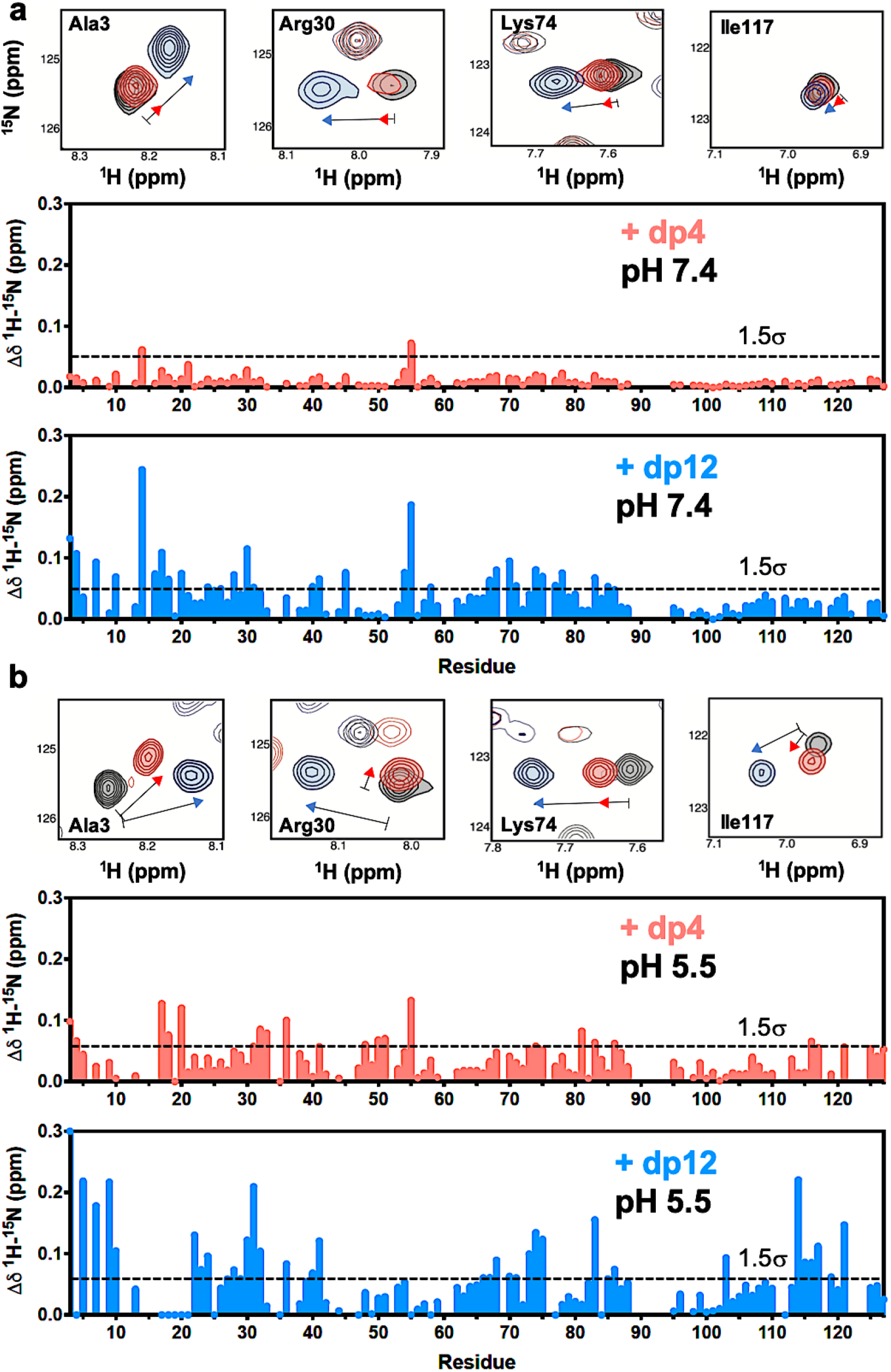
Heparin-dependent structural changes in GMCSF. **a**. Combined ^1^H^15^N chemical shift perturbations caused by dp4 (red) and dp12 (blue) binding to GMSCF at pH 7.4. Selected NMR resonances are shown to illustrate differences in chemical shift behavior of apo GMCSF (black resonances) in the presence of dp4 (red) and dp12 (blue). **b**. Combined ^1^H^15^N chemical shift perturbations caused by dp4 (red) and dp12 (blue) binding to GMSCF at pH 5.5. NMR resonances again illustrate differences in chemical shift behavior of apo GMCSF (black) in the presence of dp4 (red) and dp12 (blue). Combined chemical shift perturbations in **a** and **b** were calculated with 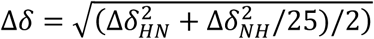. Dashed lines represent the 1.5σ above the 10% trimmed mean of all shifts measured at each pH value, and are used as a significance cut-off.

The NMR spectrum of apo GMCSF at pH 5.5 was altered more significantly by heparin oligos, in a manner distinct from that of the acidic pH itself. In the presence of unfractionated heparin at pH 5.5, the entire NMR spectrum of GMCSF is broadened beyond detection, with the exception of 10 N- and C-terminal resonances (**Figure 3b**). Presumably, this is due to the formation of a large complex with slow molecular tumbling that has been previously reported in other work.^8^Titration of pH 5.5 GMCSF with dp4 revealed 10 strongly perturbed chemical shifts, and a much greater level of overall perturbation relative to its apo GMCSF reference (**Supplementary Figure 2c**). Although the complexation of dp12 with GMCSF at pH 5.5 only induced chemical shift perturbations in 13 resonances, an additional 16 were broadened beyond detection, suggesting a change in the conformational exchange regime of these sites (**Supplementary Figure 2d**). The amino acids affected by dp12, the strongest binder of GMCSF in SPR experiments, are consistent across pH values, as 21 residues experiencing significant chemical shifts or line broadening appear in both titrations (*i*.*e*., at pH 7.4 and pH 5.5). Overall, dp12 titration caused the greatest number and magnitude of significant chemical shift perturbations at both neutral and acidic pH (**Supplementary Figure 3**). The results of these NMR titrations were mapped onto the GMCSF structure in **Figure 5**.

**Figure 5.**
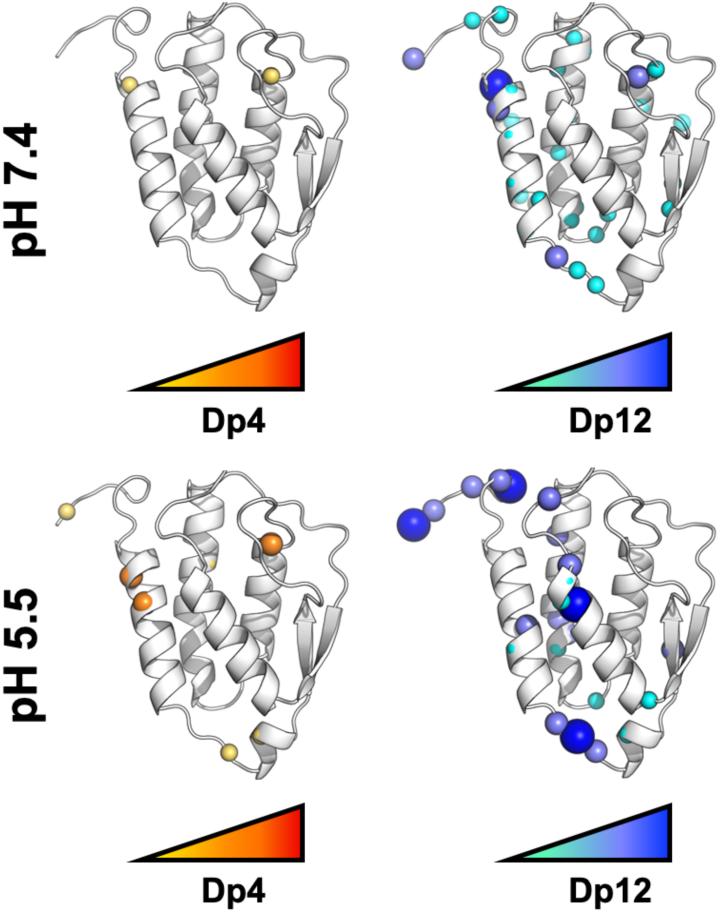
Summary of pH- and heparin size-dependent structural changes in GMCSF. Chemical shift perturbations ≥1.5σ above the 10% trimmed mean of data collected at each pH are plotted on the GMCSF structure. Spheres are sized and colored with increasing magnitude of chemical shift (dp4, yellow → red; dp12, cyan → dark blue).

### Binding of heparin with GMCSF alters its motional timescale

At neutral pH, values of the *R*_1_*R*_2_ product determined with NMR spin relaxation experiments on GMCSF-dp4 do not differ substantially from those of apo GMCSF. Saturation of GMSCF with dp12 induces µs-ms timescale motion in 8 residues (V16, L26, S44, F47, T57, M79, V116, I117, **Figure 6a**). However, when dp4 was titrated into GMCSF at pH 5.5, the dominant timescale of protein motion shifts to the ps-ns regime. Dynamics at 23 sites are clustered around the N-terminus, the linker region between helices A and β1, as well as residues within helices A and C. Only 6 residues appear flexible on the µs-ms timescale (H87, Q50, R23, S44, M79, V116). When GMCSF is titrated with dp12 at pH 5.5, 13 residues show µs-ms motion, while an additional 16 resonances are line broadened (**Figure 6b**), also suggestive of ms motion at these sites. Overall, the dynamics of GMCSF are more sensitive to the presence of heparin oligos at pH 5.5 than at pH 7.4 (**Figure 5b**). Heparin size-dependent differences in our NMR data suggest that the length of the saccharide chain affects its mode of interaction with GMCSF, and overall GMCSF behavior, when bound to each oligo.

**Figure 6.**
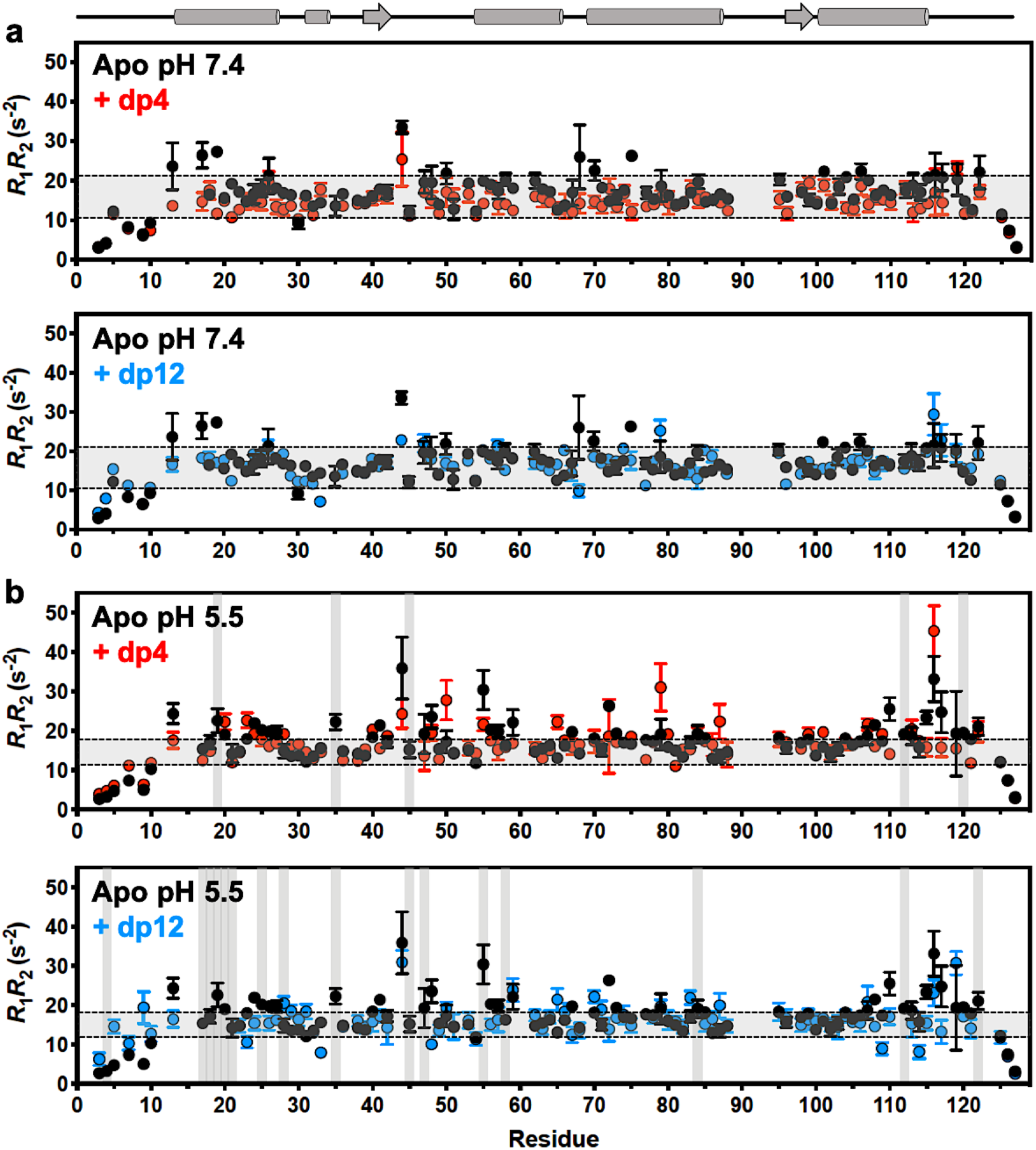
Dynamics of GMCSF-heparin complexes. **a**. Plots of *R*_1_*R*_2_ relaxation parameters determined by NMR for apo GMCSF (black circles) in the presence of dp4 (red circles) and dp12 (blue circles) at pH 7.4. Gray shaded areas mark ±1.5σ from the 10% trimmed mean of all relaxation rates determined at pH 7.4. A cartoon representation of the GMCSF structure is shown above the panel. **b**. Identical plots showing the heparin size dependence of *R*_1_*R*_2_ relaxation parameters at pH 5.5 for apo GMCSF (black) in the presence of dp4 (red) and dp12 (blue). Gray shaded areas mark ±1.5σ from the 10% trimmed mean of all relaxation rates determined at pH 5.5. Vertical gray bars represent instances of line broadening in the NMR spectrum of heparin-saturated GMCSF.

**Figure 6.**
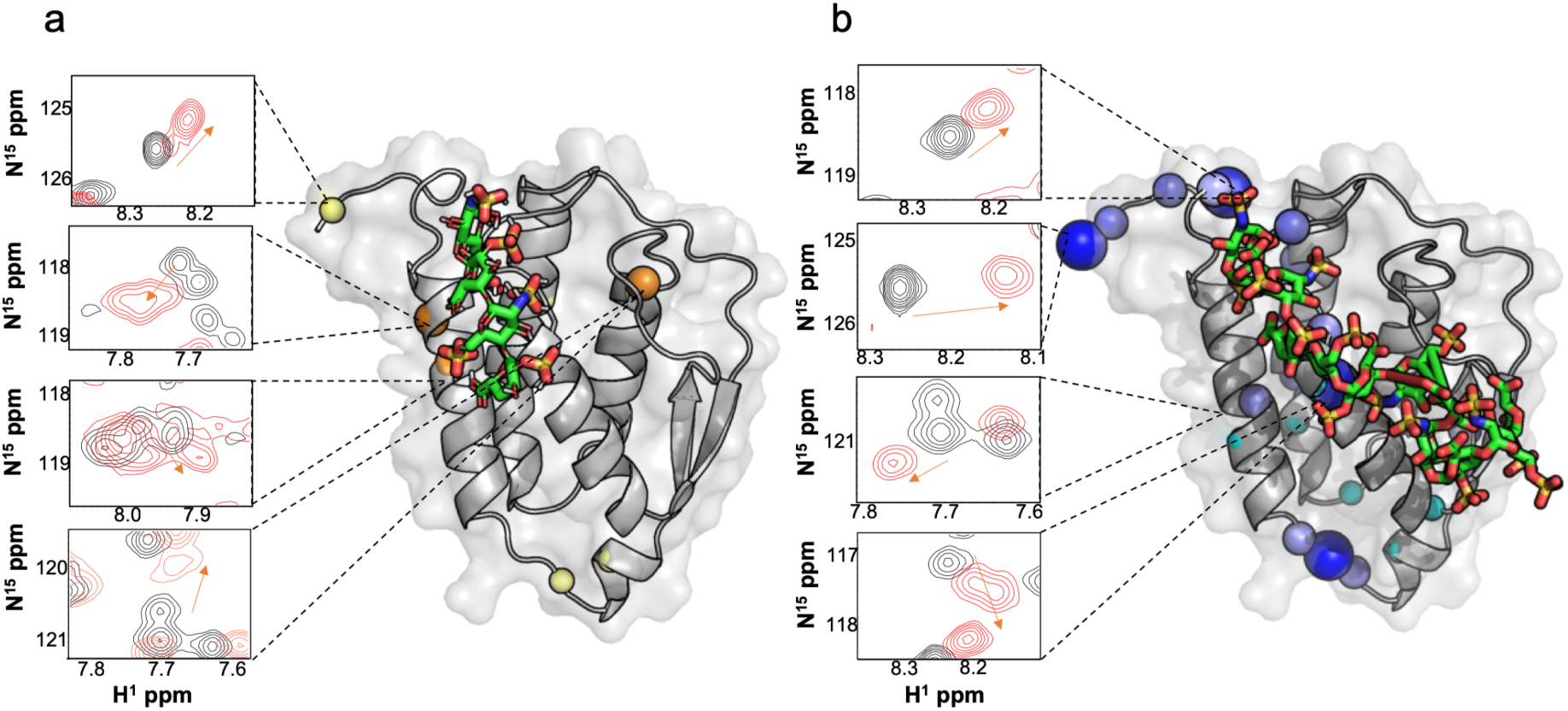
Structural models of GMCSF complexed with dp4 (**a**) and dp12 (**b**) generated with molecular docking. NMR resonances showing chemical shift perturbations from titrations of apo GMCSF (black) with dp4 or dp12 (red) are mapped along the proposed binding sites. Heparin oligos are shown as green sticks. Additional plausible structures from molecular docking are provided in the Supplementary Information.

## Discussion

GMCSF is an important mediator of innate immunity and was previously shown to form pH-dependent complexes with heparin using analytical chromatography, affinity chromatography and light scattering. ^4,7,8^ The electrostatic interaction between GMCSF and heparin is proposed to be predominantly mediated by two histidine residues, His83 and His87 that ionize at acidic pH. These residues form a positively charged binding pocket for heparin, along with Lys86. The binding interaction is abrogated upon mutation of these His residues ^7^, however, the structural changes associated with these binding interactions were not well defined. We used SPR and NMR to dissect the GMCSF-heparin interaction at the molecular level. SPR sensorgrams confirm that heparin binds to GMCSF in a pH-dependent manner. The rate of heparin association with GMCSF increases (*i*.*e*. tighter binding) at pH 4.5, and the binding affinity of heparin for GMCSF increases from *K*_*D*_ = 6.75 × 10^−6^ M to *K*_*D*_ = 1.99 × 10^−7^ M over the pH range studied (pH 7.4 - 4.5). These data are consistent with a GMCSF binding pocket that is optimized for heparin at lower pH values. Interestingly, our SPR data suggest that GMCSF and heparin interact at neutral pH as well, albeit with approximately one order of magnitude lower affinity, which has never been reported. It is likely that this interaction is transient and cannot be easily detected with biochemical assays used in previous work. Indeed, titration of heparin into GMCSF at neutral pH monitored by NMR, which readily detects both transient and tightly bound complexes, showed chemical shift perturbations consistent with an interaction. Solution NMR experiments indicate GMCSF experiences significant changes in local structure and has divergent dynamic behaviors at neutral and acidic pH. This is consistent with previously reported differences in light scattering characteristics of GMCSF at pH 7 and pH 4. ^8^ NMR experiments at pH 7.4 indicate GMCSF is generally inflexible, with the exception of freely rotating terminal residues. In contrast, at pH 5.5, a large portion of GMCSF becomes flexible on the µs-ms timescale, primarily along helix D (**Figure 1**).

It is possible that the heightened flexibility of GMCSF at pH 5.5 may facilitate stronger GMCSF-heparin contacts. We collected ^1^H^15^N HSQC spectra of apo GMCSF and with saturating amounts of unfractionated heparin, dp4 and dp12 at pH 7.4 and pH 5.5 and found that unfractionated heparin did not cause any chemical shift perturbations at pH 7.4. However, when the experiment was repeated at pH 5.5, the HSQC spectrum was almost entirely broadened beyond detection, suggesting formation of a much larger complex that limits the ability of GMCSF to tumble efficiently in solution. Although this was expected based on the average unfractionated heparin molecular weight of 15 kDa (nearly the size of GMCSF), it posed a technical challenge in that the resulting molecule appeared to be too large for NMR study, precluding useful structural information about GMCSF-heparin complexes. We, therefore, used biosynthesized heparin oligos at shorter chain lengths so that if complexes were formed with GMCSF, they would remain small enough to maintain resolution by NMR and circumvent this problem. Although it was previously reported that the molecular weight of heparin did not affect GMCSF-heparin interactions ^4^, we found that heparin oligos interact with GMCSF in a size-dependent manner (**Figures 1d, 3**). The lack of a previously observed size-dependence may be attributed to the use of heterogenous, polydisperse low molecular weight heparins, which have repeating heparin disaccharide units ranging from dp4 to dp > 24. In contrast, we found that the critical size for a difference in binding character in a homogenous population is dp > 8.

On first examination, it appears that GMCSF-heparin complexes ≥dp10 diminish sequestration of GMCSF by immobilized heparin (**Figure 1d**), thus GMCSF only efficiently binds heparins of a minimum size threshold (larger than dp8). This is also reflected in the number of residues that undergo chemical shift perturbations during NMR titrations of GMCSF with dp4 at pH 5.5, which is significantly reduced from that of the identical experiment with dp12. However, a closer examination of SPR data suggest that dp4 can increase the efficiency by which GMCSF and immobilized heparin interact compared to that of GMCSF alone. Additionally, if the change in motional timescale of flexible residues in GMCSF from µs-ms to ps-ns in the presence of dp4 but not dp12, is considered, it is possible that the binding of GMCSF with larger heparins may involve two distinct steps, only one of which is observed in experiments with smaller heparin oligos. Light scattering and sedimentation results of Wettreich and coworkers previously suggested that in order for large GMCSF-GAG complexes to form, multiple points of contact for heparin on GMCSF must exist. However, a second binding surface was not conclusively identified. Our combined SPR and NMR data suggest that GMCSF has a preferential binding site, likely mediated by strong electrostatic interactions suggested in prior work. The second binding interaction is likely made available upon the first GMCSF-heparin contact and is heparin chain length-dependent. NMR relaxation data indicate that enhanced conformational sampling, observed only in the presence of larger heparin oligos, may be crucial to optimizing additional docking points.

The primary mediators of GMCSF-heparin interactions were previously reported to be His15, His83, and His87. Interestingly, these histidine residues are not fully solvent exposed in the crystal structure of GMCSF. However, we speculate that the heightened flexibility of GMCSF at pH 5.5 may destabilize non-covalent interactions between these π-π stacked histidine residues and that protein dynamics optimize the positively charged histidine-binding pocket for negatively charged heparin. Indeed, molecular docking of dp4 onto the GMCSF structure (**Figure 6a**) demonstrates that dp4 resides in this positively charged cleft. This model is also consistent with NMR chemical shift perturbations that are localized to this site. It appears that longer heparin molecules can further interface along helices A/C, where there is another cluster of positively charged residues. These helices were found to be highly flexible in NMR relaxation experiments (**Figure 2c**) and are the sites of significant chemical shift perturbations during dp12 titrations (**Figure 6b**). These experimental signatures are exclusive to dp12, consistent with shorter heparin oligos simply not being large enough to dock with this region of GMCSF. Several other plausible models of the GMCSF-heparin complexes are provided in **Supplementary Figure 4**. These structures occupy the same general interface of GMCSF, with subtle changes in the positions of the heparin oligo chain. The binding energies of the various docking poses are comparable, and no single model provides complete coverage of the perturbations identified by NMR experiments. The pattern of chemical shifts observed during heparin oligo titrations may indicate that heparins transiently dock with GMCSF in an ensemble of these poses (**Figure 6** and **Supplementary Figure 4**) as the protein flexes on various timescales. It is unlikely that our experimental results are produced by multiple oligos binding tightly to GMCSF, as we do not observe increases in *R*_2_ values that would be sensitive to this change in molecular weight. Since these models allow limited degree-of-freedom for optimizing the ligand and protein scaffolds, they likely do not capture all of the critical interactions required for GMCSF-heparin complex formation. Thus, our future studies will be directed toward elucidating high-resolution structures of these complexes with crystallography and NMR.

## Conclusions

In this study, we probed the molecular determinants of the interaction between two clinically relevant molecules, GMCSF and heparin. Combined SPR and solution NMR studies highlight pH-sensitive binding of heparin to GMCSF that is also dependent on the length of the heparin saccharide chain. These sensitivities in the GMCSF-heparin interaction impart varying degrees of structural flexibility on GMCSF, which may be a driving force in optimizing the protein scaffold to facilitate the complex.

## Materials

Antibiotics used in protein expression and other analytical grade chemicals were purchased from Sigma. Unfractionated heparin was purchased from Sigma as a sodium salt. Heparin (16 kDa) and heparan sulfate (HS) (12 kDa from porcine intestine were purchased from Celsus Laboratories, (Cincinnati, OH). Heparin oligosaccharides including the tetrasaccharide (dp4), hexasaccharide (dp6), octasaccharide (dp8), decasaccharide (dp10), and dodecasaccharide (dp12) were prepared from controlled partial heparin lyase 1 treatment of bovine lung heparin (Sigma) followed by size fractionation.

### Protein expression and purification

Plasmid DNA containing GMCSF with an N terminal 6-His tag was cloned into a pET-15b vector and transformed into BL21(DE3) cells. GMCSF used in SPR experiments was grown and expressed in BL21(DE3) with LB medium at 37 °C. Isotopically enriched GMCSF was expressed at 37 °C in M9 minimal medium containing CaCl_2_, MgSO_4_ and MEM vitamins with ^15^NH4Cl as the sole nitrogen source. Small cultures of GMCSF were grown overnight in LB medium. The following morning, cloudy suspensions were collected by centrifugation and resuspended in the final M9 growth medium. Cultures of GMCSF were grown to an OD_600_ of 0.8-1.0 before induction with 1 mM isopropyl β-D-1-thiogalactopyranoside (IPTG).

Cells were harvested after 5 h and resuspended in a denaturing lysis buffer containing 10 mM Tris-HCl, 100 mM sodium phosphate, and 6 M guanidine hydrochloride (GuHCl) at pH 8.0. Cells were lysed by sonication and cell debris was removed by centrifugation. The resulting supernatant was incubated and nutated with 10 mL of Ni-NTA agarose beads for 30 min at room temperature before the Ni-NTA slurry was packed into a gravity column. The column was washed with the initial lysis buffer, followed by a gradient of the same buffer without GuHCl over 100 mL. Elution of GMCSF in its denatured form was performed with 1 column volume of a buffer containing 10 mM Tris-HCl, 100 mM sodium phosphate, and 250mM imidazole at pH 8.0. GMCSF was refolded by dilution via dropwise addition of the 10 mL elution into 100 mL of a refolding buffer containing 10mM Tris-HCl, 100mM sodium phosphate, and 750 mM arginine at pH 8.0. The refolded protein was dialyzed exhaustively against a buffer containing 2 mM sodium phosphate at pH 7.4. GMCSF was concentrated to ∼200 µM with an Amicon centrifugal device and stored at -20 °C.

### Surface plasmon resonance

SPR measurements were performed on a BIAcore 3000 operated using BIAcore 3000 control and BIAevaluation software (version 4.0.1). Sensor SA chips were purchased from GE healthcare (Uppsala, Sweden).

### Preparation of the heparin biochip

Biotinylated heparin was prepared by mixing 2 mg of heparin and 2 mg of amine– PEG3–Biotin (Thermo Scientific, Waltham, MA) with 10 mg of sodium cyanoborohydride (NaCNBH_3_) in 200 µL of H_2_O. The initial reaction was carried out at 70 °C for 24 h, after which an additional 10 mg of NaCNBH_3_ was added to continue the reaction for another 24 h. The mixture was then desalted with a spin column (3000 molecular weight cut-off). Biotinylated heparin was freeze-dried for chip preparation. The biotinylated heparin was immobilized on a streptavidin (SA) chip based on the manufacturer’s protocol. Briefly, a 20 μL solution of the heparin-biotin conjugate (0.1 mg/mL) in a buffer of 10 mM HEPES, 150 mM NaCl, 3 mM EDTA, and 0.005% surfactant P20 at pH 7.4 was injected over flow cell 2 (FC2), 3 (FC3) and 4 (FC4) of the SA chip at a flow rate of 10 μL/min. The successful immobilization of heparin was confirmed by the observation of a ∼100 resonance unit (RU) increase in the sensor chip signal. The control flow cell (FC1) was prepared by a 1 min injection with saturating biotin.

### Measurement of interaction between heparin and GMCSF using BIAcore

Samples of GMCSF were diluted in buffers containing 10 mM HEPES, 150 mM NaCl, 3 mM EDTA, and 0.005% surfactant P20 at pH 7.4, 5.5 and 4.5. Various dilute protein samples were injected at a flow rate of 30 µL/min. Following sample injection, the same buffer was flowed over the sensor surface to facilitate dissociation. After a 3 min dissociation time, the sensor surface was regenerated by injecting 30 µL of 0.25% sodium dodecyl sulfate (SDS). The response was monitored as a function of time (sensorgram) at 25 °C.

### Solution competition study between heparin on the chip surface and heparin-derived oligosaccharides in solution

A heparin oligosaccharide competition study was carried out by mixing 1 μM GMCSF with 1 μM of heparin oligos, including the tetrasaccharide (dp4), hexasaccharide (dp6), octasaccharide (dp8), decasaccharide (dp10), and dodecasaccharide (dp12) in 10 mM HEPES, 150 mM NaCl, 3 mM EDTA, and 0.005% surfactant P20 at pH 5.5. The mixtures were flowed over the heparin chip at a rate of 30 µL/min. After each run, the dissociation and regeneration steps were performed as described above. For each set of competition experiments, a control experiment (GMCSF without any heparin or oligosaccharides) was performed to confirm complete regeneration of the chip surface and the reproducibility of results from multiple runs (n = 2-3). Errors reported in **Table 1** were determined as standard deviations from replicate experiments.

### NMR spectroscopy

NMR samples were prepared by dialyzing 200 µM GMCSF against a buffer of 20 mM HEPES and 1 mM EDTA at pH 7.4 or pH 5.5. NMR experiments were performed on a Bruker Avance NEO 600 MHz spectrometer at 20 °C. NMR data were processed in NMRPipe and analyzed in Sparky^20,21^. NMR assignments were initially determined with triple resonance experiments at pH 7.4 and confirmed by BMRB entry 15531. Changes in resonance positions in GMCSF spectra at pH 5.5 were confirmed with standard triple resonance experiments. NMR titrations of heparins into GMCSF were performed by collecting a series of ^1^H-^15^N TROSY-HSQC spectra with increasing heparin concentration until saturation was reached (*i*.*e*., no chemical shift perturbations were observed). Trimmed means for chemical shift perturbation analyses were calculated for the entire data set (*i*.*e*. apo, +dp4, +dp12) at a given pH, in order to maintain consistency and the potential for artefacts from pooling NMR spectral data collected at different pH values. The ^1^H and ^15^N carrier frequencies were set to water resonance and 120 ppm, respectively. Longitudinal and transverse relaxation rates were determined from peak intensities of each amide resonance at multiple delay points after fitting to an exponential curve. Longitudinal and transverse relaxation rates were measured with relaxation times of 0, 20 (x2), 60 (x2), 100, 200, 600 (x2), 800, 1200, 1500, 2000, 2500 ms for *R*_1_ and 16.9, 33.9 (x2), 67.8, 136.0 (x2), 169.0, 203.0 (x2) ms for *R*_2_. Errors were determined from replicate spectra (n = 3-4). All relaxation experiments were carried out in a temperature-compensated interleaved manner, were processed with in-house scripts, and analyzed in GraphPad Prism 7.0 (GraphPad Software).

### Computational methods

Automated docking simulations were conducted with AutoDock 4.2 (Scripps Research Institute, LaJolla, CA) using crystal structure PDB:2GMF ^22^ to generate atomic energy grids ^23^. Heparin oligo structures dp4 and dp12 were generated as ligand files using custom PDB files as annotated in the AutoDock User Guide. All water molecules were removed, hydrogen atoms were added to the crystal structure and atomic partial charges were assigned to the protein atoms within the AutoDock software. The Lamarckian genetic algorithm was used while a 100Å cubic box centered on His83 with 0.5 Å grid positions was used. All histidine residues were set to be protonated to emulate the low pH state of GMCSF. Docking studies were run for both dp4 and dp12 heparin oligos. Results were retrieved as a collection of poses representative of regions on GMCSF occupied by heparin.

### Data availability

Plasmid DNA used in this work is freely available upon request. NMR pulse sequences used in this work were directly from the Bruker Pulse Program library. NMR processing scripts are freely available upon request.

## Acknowledgments

This work was supported by start-up funds from Brown University (to GPL).

## Author Contributions

J.Y.C. expressed and purified GMCSF, collected and analyzed NMR and computational data; F.Z. and R.J.L. produced heparin oligos, collected and analyzed SPR data; G.P.L. supervised the project and analyzed NMR data. The manuscript was written through contributions of all authors.

## Additional Information

Supplementary Information is available for this paper

## Protein Accession Information

Granulocyte macrophage colony-stimulating factor (*homo sapiens*) – UniProtKB P04141

